# Reduction in cardiolipin reduces expression of creatine transporter-1 and creatine transport in hBMEC/D3 human brain microvessel endothelial cells

**DOI:** 10.1101/2022.01.29.478231

**Authors:** Donald W. Miller, Grant M. Hatch

## Abstract

**Background:** Cardiolipin (CL) is the signature phospholipid of mitochondria and regulates a plethora of cellular functions including mitochondrial energy production. There is little knowledge of how CL regulates uptake and membrane transport processes in mammalian cells. Endothelial cells of the blood brain barrier (BBB) play a vital role in uptake of metabolites into the brain and are enriched in mitochondria compared with peripheral endothelial cells. We examined how deficiency in BBB endothelial CL regulates the expression of selected drug and metabolite transporters and their function.

**Methods:** Cardiolipin synthase-1 (hCLS1) was knocked down in a human brain microvessel endothelial cell line, hBMEC/D3, by Lipofectamine^®^ transfection with hCLS1 siRNA and CL levels, CL synthase activity and the mRNA expression of selected blood brain barrier drug and metabolite transporters examined. Mock transfected hBMEC/D3 cells served as controls. The incorporation of [^14^C]creatine and [^14^C]oleate into hBMEC/D3 cells was determined as a measure of solute metabolite transport and protein expression of the creatine transporter determined.

**Results:** Knockdown of the CL biosynthetic enzyme hCLS1 in hBMEC/D3 reduced CL and CL synthase activity and the mRNA expression of creatine transporter-1, p-glycoprotein and breast cancer resistance protein compared to controls. In contrast, mRNA expression of ATP binding cassette subfamily C members-1, -3, multidrug resistance-associated protein-4 variants 1, -2, and fatty acid transport protein-1 were unaltered. Although ATP production was unaltered by hCLS1 knockdown, incorporation of [^14^C]creatine into hBMEC/D3 cells was reduced compared to controls. The reduction in [^14^C]creatine incorporation was associated with a reduction in creatine transporter-1 protein expression. In contrast, incorporation of [^14^C]oleic acid into hBMEC/D3 cells and the mRNA expression of fatty acid transport protein-1 was unaltered by knockdown of hCLS1 compared to controls.

**Conclusion:** Knockdown of hCLS1 in hBMEC/D3, with a corresponding reduction in CL, results in alteration in expression of specific solute membrane transporters.

## Introduction

Cardiolipin (CL) is a major polyglycerophospholipid in mammalian tissues and comprises approximately 7-15% of the entire phospholipid phosphorus mass of mitochondria^1-3^. CL modulates the activity of many mitochondrial enzymes involved in the generation of ATP (reviewed in^3,4^) and is the “glue” that holds the mitochondrial electron transport chain together^5,6^. However, CL has recently been implicated in the regulation of many other cellular processes including apoptosis, autophagy, mitophagy and membrane transport of glucose^7-10^. The final step of *de novo* biosynthesis of CL is the conversion of phosphatidylglycerol to CL catalyzed by CL synthase (CLS) localized exclusively to the inner mitochondrial membrane^11,12^. We previously cloned and characterized the murine and human CLS (hCLS1)^13^.

BTHS is a rare X-linked genetic disorder, caused by a mutation in the TAFAZZIN gene, and is the only disease in which the specific biochemical defect is a reduction in CL (reviewed in^14,15^). Previous studies have suggested that some BTHS patients exhibit a cognitive phenotype^16^. In addition, abnormal CL levels have also been reported in numerous neurodegenerative disorders (reviewed in^7^). We recently examined how tafazzin deficiency impacts cognitive function in mice^17^. We showed that tafazzin knockdown in mice resulted in significant memory deficiency based on novel object recognition test. Whether the cognitive defects in tafazzin knockdown mice are related to a loss in CL in blood brain barrier (BBB) endothelial cells and/or transport of compounds into the brain is unknown. In fact, few studies have examined if altering CL levels regulate uptake and membrane transport processes in mammalian BBB endothelial cells.

In the current study we examined if reduction in CL altered the expression and function of selected BBB endothelial cell transport proteins. We show that knockdown of hCLS1 in hBMEC/D3 human BBB endothelial cells, with a corresponding reduction in CL, results in reduced expression of creatine transporter-1 expression and a reduction in [^14^C]creatine incorporation into these cells.

## Materials and Methods

### Materials

Cell culture medium (CSC complete medium) and passaging reagents were obtained from Cell Systems Corporation (Kirkland, WA, USA). Human adult brain endothelial cell line hCMEC/D3 was kindly provided by Dr. P.O. Couraud (Institut Cochin, France). Endothelial basal medium-2 was from Lonza (Basal, Switzerland). Fetal bovine serum (FBS), other medium supplements, cell culturing reagents and primers and reagents used for qPCR were obtained from Life Technologies Inc. (Burlington, ON, Canada) and Sigma-Aldrich (St. Louis, MO, USA). Opti-MEM reduced serum medium, Lipofectamine RNAiMAX Transfection Reagent and the Silencer were obtained from Thermo Fisher (Winnipeg, MB, Canada). Select Pre-designed siRNA for human cardiolipin synthase-1 (CLS) were obtained from Life Technologies Inc. (Burlington, ON, Canada). RNeasy and Plus Mini Kit used for RNA extraction was obtained from Qiagen (Cambridge, MA, USA). Silencer® Select Negative Control No.2 siRNA was from Life Technologies (Burlington, ON). [1-14C]oleate and [^14^C]creatine were obtained from PerkinElmer (Boston, MA, USA). Ecolite scintillant was obtained from ICN Biochemicals (Montreal, QC, Canada). Seahorse XF24 analyzer and reagent kits for determination of the relative contribution of glycolysis and oxidative phosphorylation (OX-PHOS) to basal ATP production were from Aligent (North Billerica, MA, USA). Antibodies to SLC6A8 (Crt1) and cyclophilin were from ThermoFisher (Winnipeg, Canada). HRP-linked monkey anti-rabbit secondary antibody was from GE Healthcare Life Sciences (Little Chalfont, UK). All other biochemicals were of ACS grade and were obtained from either Sigma-Aldrich or Fisher Scientific (Winnipeg, MB, Canada).

### Transfection of cells

HCMEC/D3 cells were transfected for 48 h with human cardiolipin synthase-1 (hCLS1) siRNA to knock down the CL *de novo* biosynthetic enzyme hCLS1 as previously described^10^. In brief, transfection of cells was performed using siRNA against the hCLS1 gene and lipofectamine^®^ RNAiMAX transfection reagent kit according to the manufacturers protocol. The siRNA target sequence for hCLS1 was 5’-GGACAAUCCCGAAUAUGUUtt-3’. The Silencer^®^ Select Negative Control No.2 siRNA was used in the mock transfection as a control. It has been tested using microarray analysis and shown to have minimal effects on gene expression profiles.

### mRNA expression analysis

hCMEC/D3 cells were grown and transfected in 100 mm cell culture dishes. After 48 h of transfection, cells were harvested and total RNA was obtained using the RNeasy® Plus Mini kit and Qiashredder homogenizer columns (Qiagen). The integrity of total RNA was confirmed by running the RNA sample on a denaturing agarose gel. Gene expression analysis was measured using the Mastercycler ep *realplex* system (Eppendorf). Primers were designed using NCBI/Primer-Blast and were synthesized by Invitrogen (Ontario, Canada). The primers and annealing temperatures used for RT-PCR analysis are outlined in **Table 1**. In addition to fatty acid transport protein-1 (FATP-1) and the creatine transporter (Crt1), mRNA expression of the drug efflux transporters P-glycoprotein (P-gp), breast cancer resistance protein (BCRP), multidrug resistance-associated protein-4 variants (MRP4_v1, MRP4_v2) and ATP binding cassette subfamily C members-1, -3 (ABCC1, ABCC3) in hCMEC/D3 cells were determined since these transporters are expected to be found in brain endothelial cells^18^. Expression was normalized to 18S RNA.

**Table 1.**
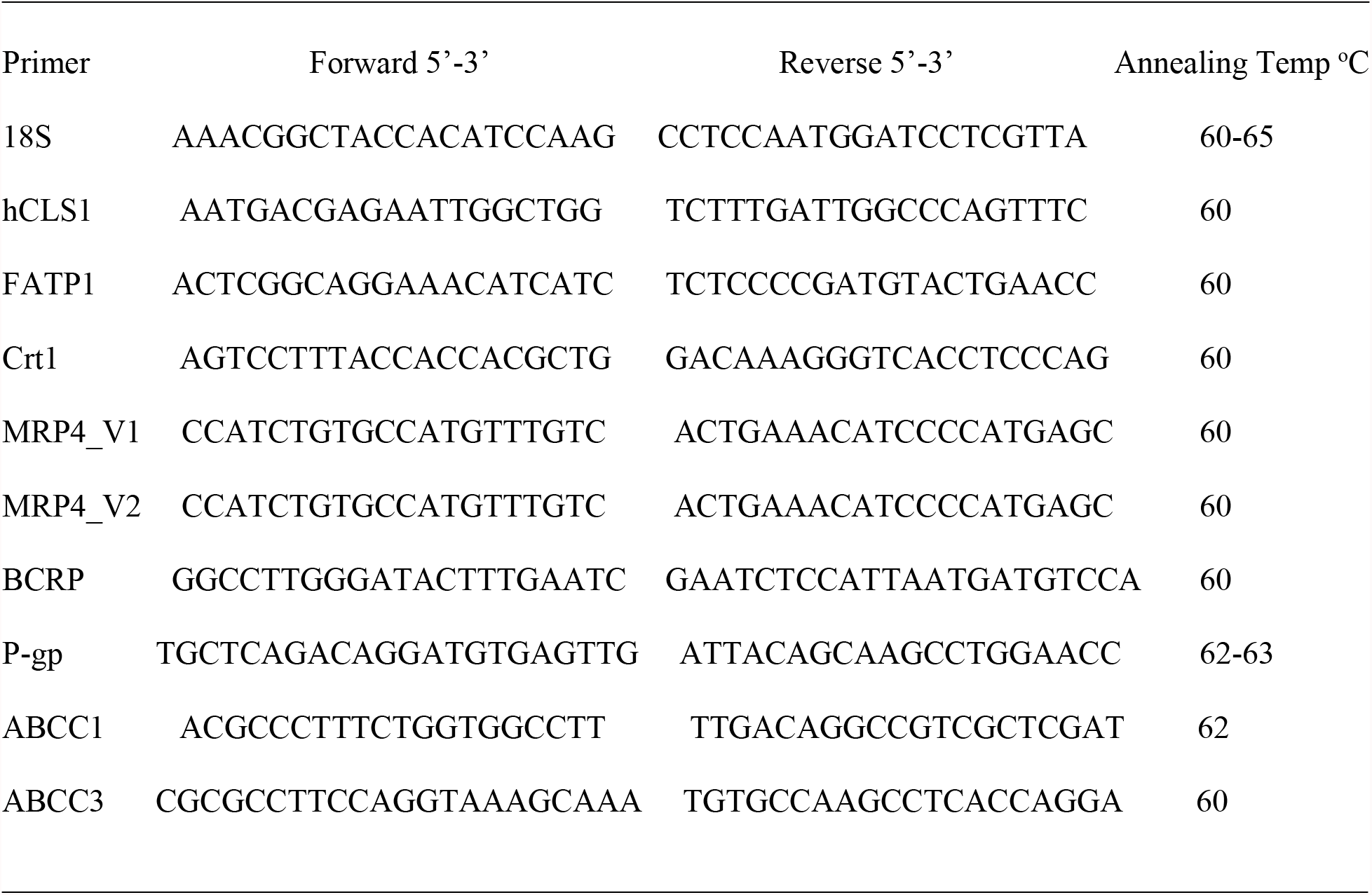
Primers and annealing temperatures used for RT-PCR.

### Radiolabeling of cells

Incorporation of radiolabeled creatine (as a measure of creatine transporter dependent solute transport) and uptake of radiolabeled oleate (as a measure of fatty acid transport) into cells was performed. Control and hCLS1 siRNA transfected cells were incubated with 0.1 mM [1-^14^C]oleate (bound to albumin, 1:1 molar ratio) for up to 30 min or with [^14^C]creatine for 10 min and radioactivity incorporated into cells determined as described^10^.

### Western blot analysis and determination of CL levels

Equal amounts of mitochondrial proteins (40 µg/lane) were loaded and separated by electrophoresis on a 12% SDS-PAGE gel. The proteins were then transferred onto a PVDF transfer membrane (Immobilon, Millipore, Bedford, MA). The presence of transferred proteins on the membrane was confirmed by staining with Ponceau S (Sigma). Membranes were blocked for 2 hours at room temperature with 5% non-fat milk in 0.1% tween-20/TBS (TBS-T). Then, membranes were incubated overnight at 4°C in blocking buffers with rabbit primary antibodies against Crt1 (1:150 dilution) or cyclophilin. Expression of cyclophilin was used as the loading control. After several washes with TBS-T, membrane was incubated with HRP-linked monkey anti-rabbit secondary antibody (1:5000) at room temperature for 1 hour. Protein bands in the membranes were then visualized by enhanced chemiluminescence. The relative intensities of the bands were analyzed by Image J software and normalized to cyclophilin.

Lipids were extracted from cells and CL was separated using thin layer chromatography as previously described^10^. The plates were stained with iodine and spots corresponding to CL removed and lipid phosphorus determined as described^19^.

### Relative contribution of glycolysis and oxidative phosphorylation to the basal ATP production rate determination

The oligomycin sensitive oxygen consumption rate (OCR) and the glycolytic proton production rate (both measured under a saturating extracellular glucose concentration) were determined using a Seahorse XF24 analyzer with reagent kits as per the manufacturer’s instructions and converted to ATP production rate. The proton production rate expressed as pmol H+/min was automatically converted by the Seahorse XF-24 analyzer from the extracellular acidification rate (mpH/min) using the buffer capacity of the media and the chamber volume. The oligomycin sensitive OCR was converted to ATP production using a P/O ratio of 2.3, while the glycolytic PPR was converted to ATP production using a 1:1 ratio, based on the fact that in glycolysis one ATP is made per lactate/proton produced^20^.

### Statistical analysis

All data were expressed as mean ± S.D. The differences between the experimental and the control groups were evaluated by Student’s t-test. All values with p < 0.05 were considered statistically significant.

## Results

Transfection of hCMEC/D3 cells with human cardiolipin synthase-1 (hCLS1) siRNA reduced hCLS1 levels approximately 70% compared to mock-transfected control cells (**Fig. 1A**). This resulted in an approximate 60% reduction in CLS enzyme activity in hCLS1 siRNA transfected cells compared to mock-transfected controls (**Fig. 1B**). In addition, CL levels were reduced 35% (n=3-4, p<0.05) from 3.5±0.6 pmol/mg protein to 2.3±0.5 pmol/min/mg protein in hCLS1 siRNA transfected cells compared to mock-transfected controls, respectively. Thus, knockdown of hCLS1 in hBMEC/D3 cells reduces CL levels.

**Figure 1.**
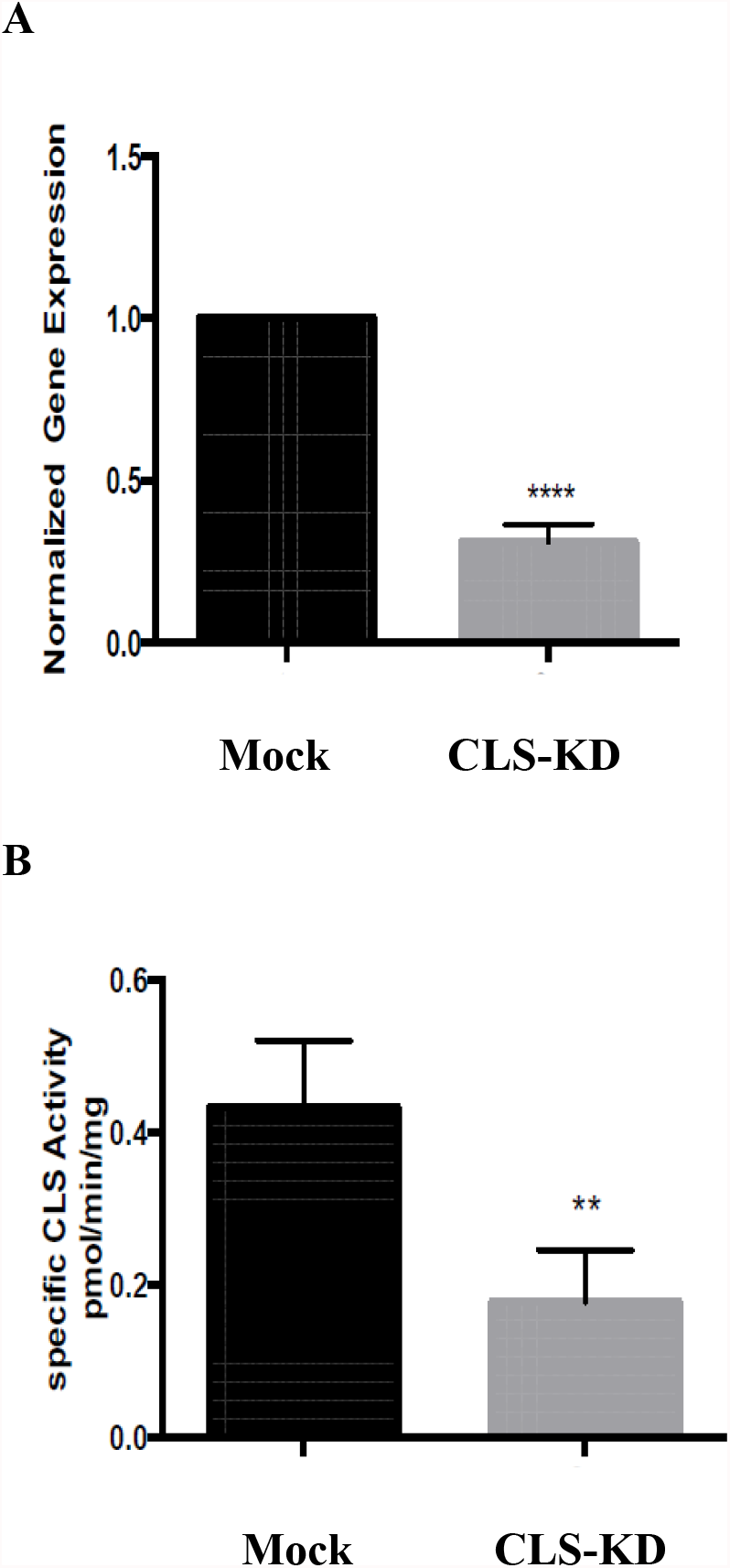
hCLS1 mRNA expression and CLS enzyme activity in hBMEC/D3 cells transfected with hCLS1 siRNA. hBMEC/D3 cells were mock transfected or transfected with hCLS1 siRNA for 48 h and hCLS1 mRNA expression (**A**) and CLS enzyme activity (**B**) determined as described in Materials and Methods. n=3-4, **p<0.05; ****p<0.001.

We initially examined mRNA expression of the fatty acid transport protein FATP1 and the multidrug resistance-associated protein-4 variants MRP4_v1 and MRP4_v2 in hCMEC/D3 CLS-KD cells. mRNA expression of FATP1, MRP4_v1 and MRP4_v2 were unaltered in hCLS1 siRNA transfected cells compared to mock-transfected controls (**Fig. 2A**). We previously determined that membrane integrity was maintained in hCMEC/D3 cells with hCLS1 siRNA knockdown^4^. This was confirmed by examining the incorporation of [1-^14^C]oleate into hCMEC/D3 cells. Incorporation of [1-^14^C]oleate into hCMEC/D3 cells was identical between mock-transfected controls and hCLS1 siRNA transfected cells incubated under low (1 mM) glucose conditions and corresponded to the lack of alteration in FATP1 mRNA expression in these cells (**Fig. 2A,B**). Membrane integrity can be potentially compromised by loss of ATP. We confirmed that hCLS1 knock down did not affect cellular ATP production rate (**Fig. 2C**).

**Figure 2.**
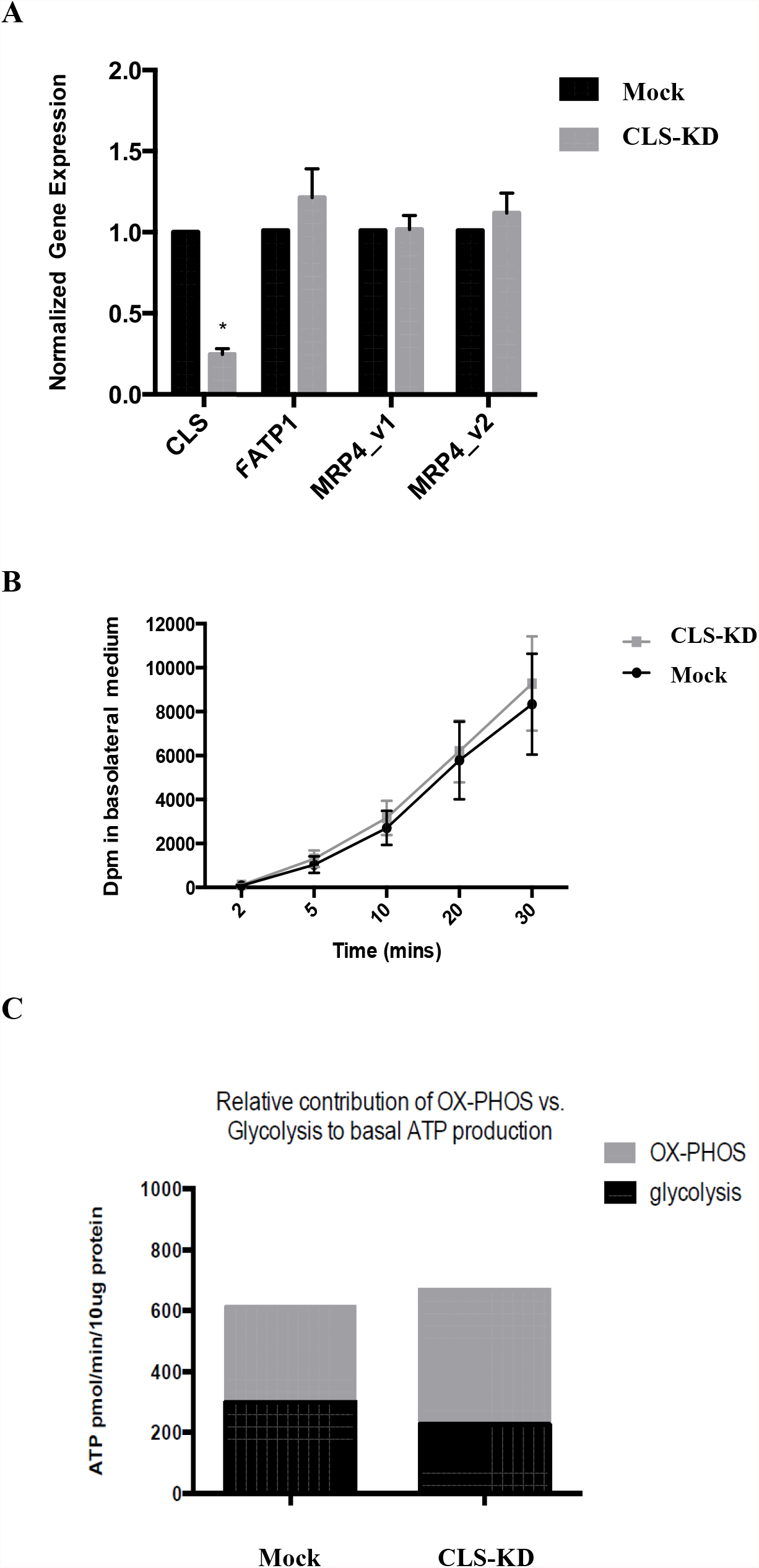
Expression of FATP1, MRP4_v1 and MRP4_v2, [1-^14^C]oleate uptake and relative contribution of oxidative phosphorylation and glycolysis to ATP production in hBMEC/D3 cells transfected with hCLS1 siRNA. hBMEC/D3 cells were mock transfected or transfected with hCLS1 siRNA for 48 h and FATP1, MRP4_v1 and MRP4_v2 mRNA expression (**A**), [1-^14^C]oleate uptake (**B**) and relative contribution of oxidative phosphorylation (OX-PHOS) and glycolysis to ATP production (**C**) determined as described in Materials and Methods. n=3-4, *p<0.05.

We next examined mRNA expression of other major transporters in hBMEC/D3 CLS-KD cells that are expected to be found in brain endothelial cells^18^. mRNA expression of Crt1, P-gp, and BCRP were reduced by hCLS1 knockdown but not the mRNA expression of ABCC1 and ABCC3 (**Fig. 3A**). The greatest reduction was in Crt1 (45% decrease, p<0.05). In addition, protein expression of Crt1 and [^14^C]creatine uptake into hBMEC/D3 cells were reduced by hCLS1 knockdown (**Fig. 3B, 3C**). Thus, knockdown of hCLS1 in hBMEC/D3, with a corresponding reduction in CL, results in reduction in Crt1 expression and [^14^C]creatine uptake into hBMEC/D3 cells.

**Figure 3.**
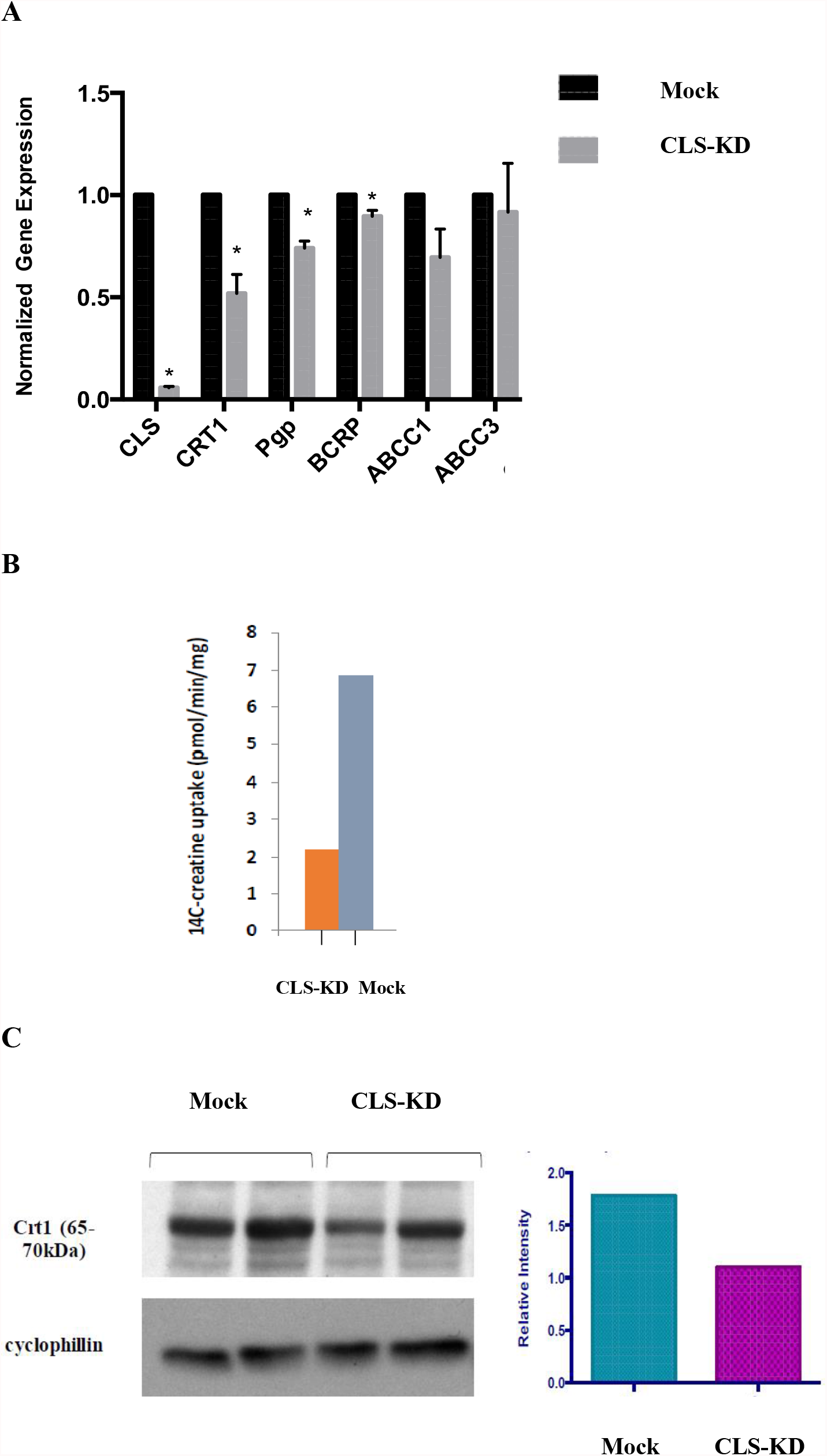
Expression of Crt1, P-gp, BCRP, ABCC1 and ABCC3, and [^14^C]creatine uptake, and Crt1 protein expression in hBMEC/D3 cells transfected with hCLS1 siRNA. hBMEC/D3 cells were mock transfected or transfected with hCLS1 siRNA for 48 h and Crt1, P-gp, BCRP, ABCC1 and ABCC3 mRNA expression (**A**), [^14^C]creatine uptake (**B**) and Crt1 protein expression (**C**) determined as described in Materials and Methods. n=3-4, *p<0.05.

## Discussion

In the current study we examined if deficiency in BBB endothelial CL in hBMEC/D3 cells regulate the expression of selected drug and metabolite transporters. We observed that knockdown of hCLS1 in hBMEC/D3 cells reduced CLS activity and CL levels and the mRNA expression of Crt1, P-gp and BCRP, but not other transporters examined. The reduction in Crt1 mRNA expression resulted in a corresponding reduction in Crt1 protein expression and a lowered incorporation of [^14^C]creatine into hBMEC/D3 cells. These data suggest that optimal CL levels are required for the regulation of creatine uptake into hBMEC/D3 cells.

Three families of ABC transporters have been predominantly characterized to play important role in the transport of therapeutic drugs across the BBB including P-glycoprotein (P-gp), the breast cancer resistance protein (BCRP), and the multi-drug resistance associated proteins (MRPs)^21^. P-gp is localized on the abluminal surface of BBB endothelial cells and functions in preventing many lipophilic substances from entering into the brain. A wide range of pharmaceutical agents with molecular weights ranging from 3000-4000 Da are reported to be substrates of this transporter. These compounds include drugs such as antibiotics, immunosuppressive agents and anti-cancer agents^21,22^. Interestingly, P-gp expression was reported to be increased in neurological conditions such as brain tumors and epilepsy and may partly be responsible for the multi-drug resistance observed in these conditions. Thus, strategies to either inhibit P-gp or down-regulate its expression would be advantageous in the treatment of these neurological disorders. Indeed, many P-gp inhibitors have been developed and are in different stages of clinical trials^21^. However, most of these inhibitors have failed to be clinically successful. This may, in part, be due to the fact that sufficient expression of P-gp is required under physiological conditions to keep a wide range of neurotoxins from accumulating in the brain. Indeed, several studies have demonstrated that normally well tolerated drugs become neurotoxic in the absence of P-gp due to high accumulation in the brain (reviewed in^23^). BCRP was initially discovered in a multi-drug resistant breast cancer cell line that exhibited reduction in accumulation of chemotherapeutic drugs, even in the absence of P-gp and MRPs^24^. Due to its sequence and structural homology, BCRP was suggested to belong to the ABCG family, a subfamily of the ABC superfamily of transporters^25^. BCRP expression is not specific to chemotherapy resistant breast cancer cells^24^. Since high expression of P-gp mediates resistance of cancers to chemotherapeutic drugs, our findings of reduced P-gp and BCRP mRNA expression in hCLS1 knockdown hBMEC/D3 cells indicate that lowering CL levels in BBB endothelial cells could potentially be used as a therapeutic approach to enhance drug entry into the brain to treat brain tumors.

MRPs are members of the ABC superfamily known to play an important role in multidrug resistance. MRP family members transport mainly anionic drugs such as methotrexate but their substrates may also include neutral agents such as glutathione and its derivatives^26^. MRPs substrate specificity may overlap with P-gp to further enhance drug resistance. We observed that, in addition to FATP1, reduction in CL in hBMEC/D3 cells did not affect mRNA expression of the multidrug resistance-associated protein-4 variants MRP4_v1 and MRP4_v2 or ABCC1 and ABCC3 indicating that loss of CL impact only selected transporters in hBMEC/D3 cells.

In organs that require a high burst of energy supply during activation such as the muscle and brain, ATP is often stored in the form of phosphocreatine that can serve as an immediate substrate for ATP regeneration - a process 12 times faster than oxidative phosphorylation (OX-PHOS)^27^. Indeed, this is supported by the observation that phosphocreatine levels rapidly decrease during brain activation while the ATP levels remain constant. Whether creatine is primarily synthesized in brain or is imported from the peripheral circulation remains a subject of debate. Crt1 is encoded by the SLC6A8 gene and exhibits the ability to uptake creatine into the brain^28,29^. However, as expression of this transporter is absent in the astrocytic end feet covering the endothelium, it was suggested that the permeability of creatine into the brain might be limited^29^. This may indeed be true as oral administration of 20 g of creatine per day consecutively for 4 weeks in healthy volunteers only resulted in a moderate 5-10% increase in brain creatine levels^30^. As the brain is also known to express both L-arginine:glycine amidinotransferase and guanidinoacetate methyltransferase, enzymes required for endogenous creatine synthesis, it has been proposed that the brain can self-supply most of its required creatine^29^. This is supported by the fact that treatment of L-arginine:glycine amidinotransferase and guanidinoacetate methyltransferase deficient patients with a high oral doses of creatine partially replenished cerebral creatine levels but did not impact these patients central nervous system developmental alterations^31^. These observations challenge the importance of Crt1 in supplying creatine to the brain. However, the fact that patients with defects in the SLC6A8 gene experience progressive brain atrophy, cognitive disability and speech deficit which cannot be treated by creatine oral supplementation, indicate an essential role of Crt1 in maintaining healthy brain creatine levels. In the current study, the reduction in uptake of creatine was not likely due to reduction in cellular ATP due to reduced CL since ATP production rate appeared to be maintained in cells with knockdown of hCLS1. This is in agreement with our previous study in which CLS knockdown with reduced CL resulted in an increased glucose uptake in order to maintain cellular ATP levels^10^. Interestingly, control hCMEC/D3 cells appeared to make ∼60% of their ATP via OX-PHOS under basal condition. This supports the hypothesis that, unlike endothelial cells in non-BBB regions, brain capillary endothelial cells may be more dependent on OX-PHOS for their energy requirements under basal conditions. A limitation of our study is that it does not address whether the reduction in CL plays a direct or indirect role on membrane transport. However, the observation that FATP1 mRNA expression and [1-^14^C]oleate uptake, a process dependent on energy, are unaltered in hBMEC/D3 cells with knockdown of CLS coupled with a normal ATP production rate in these cells clearly indicate that the effect of reduced CL on [^14^C]creatine transport is not simply due to alteration in energy metabolism.

## Acknowledgements

We thank Hieu M. Nguyen, Ngoc On and Fred Y. Xu for performing experiments. This work was supported by an NSERC grant to GMH and CIHR grant to DWM. GMH is the Canada Research Chair in Molecular Cardiolipin Metabolism.

## References

1. Pangborn M. (1942) J. Biol. Chem. 143, 247.

2. Poorthuis BJHM, Yazaki PJ, Hostetler KY. (1976) J. Lipid Res. 17, 433.

3. Hatch GM. (2004) Biochem. Cell Biol. 82, 99.

4. Hoch FL. (1992) Biochim. Biophys. Acta 1113, 71.

5. Zhang M, Mileykovskaya E, Dowhan W. (2002) J. Biol. Chem. 277, 43553.

6. Ascenzi P, Polticelli F, Marino M, Santucci R, Coletta M. (2011) IUBMB Life. 63, 160.

7. Falabella M, Vernon HJ, Hanna MG, Claypool SM, Pitceathly RDS. (2021) Trends Endocrinol. Metab. 32, 224.

8. Hsu P, Shi Y. (2017) Biochim Biophys Acta Mol Cell Biol Lipids. 1862, 114.

9. Choubey V, Zeb A, Kaasik A. (2021) Cells. 11, 38.

10. Nguyen HM, Mejia EM, Chang W, Wang Y, Watson E, On N, Miller DW, Hatch GM. (2016) J. Neurochem. 139, 68.

11. Hostetler KY, Van den Bosch H. (1971) Biochim. Biophys. Acta 239, 113.

12. Schlame M, Haldar D. (1993) J. Biol. Chem. 268, 74.

13. Lu B, Xu F, Jiang J, Choy PC, Hatch GM, Grunfel C, Feingold K. (2006) J. Lipid Res. 47, 1140.

14. Hauff K, Hatch GM. (2006) Prog. Lipid Res. 45, 91.

15. Barth PG, Valianpour F, Bowen VM, Lam J, Duran M, Vaz FM, Wanders RJ. (2004) Am. J. Med. Genet. 126A, 349.

16. Mazzocco MM, Henry AE, sKelly RI (2007) J. Dev. Behav. Pediatr. 28, 22–30.

17. Cole LK, Kim JH, Amoscato AA, Tyurina YY, Bayir H, Kagan V, Hatch GM, Kauppinen TM (2018) Biochim. Biophys. Acta - Molec. Basis Dis. 1864, 3353.

18. Dalvi S, On N, Nguyen H, Pogorzelec M, Miller DW, Hatch GM (2014) In: Neurochemistry (Heinbockel, T., ed.) http://www.intechopen.com/books/neurochemistry/the-blood-brain-barrier-regulation-of-fatty-acid-and-drug-transport. In Tech, pp. 1.

19. Rouser G, Fkeischer S, Yamamoto A. (1970) Lipids 5: 494.

20. Brand MD (2005) Biochem. Soc. Trans. 33, 897.

21. Loscher W, Potschka H. (2005) NeuroRx 1, 86.

22. Miller DS, Bauer B, Hartz AM. (2008) Pharmacol. Rev. 60, 196.

23. Schinkel AH. Adv. Drug Deliv. Rev. (1999) 36, 179.

24. Doyle LA, Yang W, Abruzzo LV, Krogmann T, Gao Y, Rishi AK, Ross DD. (1998) Proc. Natl. Acad. Sci. U.S.A. 95, 15665.

25. Schinkel AH, Jonker JW. (2003) Adv. Drug Deliv. Rev. 55, 3.

26. Borst P, Evers R, Kool M, Wijnholds J. (2000) J. Natl. Cancer Inst. 92, 1295.

27. Rae C, Digney AL, McEwan SR, Bates TC. (2003) Proc. Biol. Sci. 270, 2147.

28. Ohtsuki S, Tachikawa M, Takanaga H, Shimizu H, Watanabe M, Hosoya K, Terasaki T. (2002) J. Cereb. Blood Flow Metab. 22, 1327.

29. Beard E, Braissant O. (2010) J. Neurochem. 115, 297.

30. Dechent P, Pouwels PJ, Wilken B, Hanefeld F, Frahm J. (1999) Am. J. Physiol. 277, R698.

31. Braissant O. (2012) J. Inherit. Metab. Dis. (2012) 35, 655.

